# The International Weed Genomics Consortium: Community Resources for Weed Genomics Research

**DOI:** 10.1101/2023.07.19.549613

**Authors:** Jacob S. Montgomery, Sarah Morran, Dana R. MacGregor, J. Scott McElroy, Paul Neve, Célia Neto, Martin M. Vila-Aiub, Maria Victoria Sandoval, Analia I. Menéndez, Julia M. Kreiner, Longjiang Fan, Ana L. Caicedo, Peter J. Maughan, Bianca Assis Barbosa Martins, Jagoda Mika, Alberto Collavo, Aldo Merotto, Nithya K. Subramanian, Muthukumar V. Bagavathiannan, Luan Cutti, Md. Mazharul Islam, Bikram S Gill, Robert Cicchillo, Roger Gast, Neeta Soni, Terry R. Wright, Gina Zastrow-Hayes, Gregory May, Jenna M. Malone, Deepmala Sehgal, Shiv Shankhar Kaundun, Richard P. Dale, Barend Juan Vorster, Bodo Peters, Jens Lerchl, Patrick J. Tranel, Roland Beffa, Alexandre Fournier-Level, Mithila Jugulam, Kevin Fengler, Victor Llaca, Eric L. Patterson, Todd Gaines

## Abstract

The International Weed Genomics Consortium is a collaborative group of researchers focused on developing genomic resources for the study of weedy plants. Weeds are attractive systems for basic and applied research due to their impacts on agricultural systems and capacity to swiftly adapt in response to anthropogenic selection pressures. Our goal is to use genomic information to develop sustainable and effective weed control methods and to provide insights about biotic and abiotic stress tolerance to assist crop breeding. Here, we outline resources under development by the consortium and highlight areas of research that will be impacted by these enabling resources.

## Introduction

Each year globally, agricultural producers and landscape managers spend billions of US dollars [1, 2] and countless hours attempting to control weedy plants and reduce their adverse effects. These management methods range from low-tech (e.g., pulling plants from the soil by hand) to extremely high-tech (e.g., computer vision-controlled spraying of herbicides). Regardless of technology level, effective control methods serve as strong selection pressures on weedy plants, and often result in rapid evolution of weed populations resistant to such methods [3–7]. Thus, humans and weeds have been locked in an arms race, where humans develop new or improved control methods and weeds adapt and evolve to circumvent such methods.

Applying genomics to weed science will enable the development of more sustainable and effective control methods and offer a unique opportunity to study rapid adaptation and evolutionary rescue of diverse weedy species in the face of widespread and powerful selective pressures. Furthermore, lessons learned from these studies may also help to improve crop breeding efforts in the face of our ever-changing climate. While other research fields have used genetics and genomics to uncover the basis of many biological traits [8–11] and to understand how ecological factors affect evolution [12, 13], the field of weed science has lagged behind in the development of genomic tools essential for such studies [14]. As research in human and crop genetics pushes into the era of pangenomics, (i.e., multiple chromosome scale genome assemblies for a single species [15, 16]) publicly available genomic information is still lacking or severely limited for the majority of weed species. In fact, a recent review of current weed genomes identified just 26 weed species with sequenced genomes [17] – many assembled to a sub-chromosome level.

The International Weed Genomics Consortium (IWGC) is an open collaboration between academic, government, and industry researchers focused on producing genomic tools for weedy species from around the world. Through this collaboration, our initial aim is to provide chromosome-level reference genome assemblies for at least 50 important weedy species from across the globe. Each genome will include annotation of gene models and repetitive elements and will be freely available through public databases with no intellectual property restrictions. Species were chosen based on member input, economic impact, and global prevalence (Figure 1). Additionally, future funding of the IWGC will focus on supplementing these reference genomes with tools that increase their utility.

**Figure 1.** The International Weed Genomics Consortium (IWGC) collected input from the weed genomics community to develop plans for weed genome sequencing, annotation, user-friendly genome analysis tools, and community engagement.

The IWGC held its first conference in Kansas City, Missouri, USA in September of 2021. At this meeting, guest speakers highlighted successful examples of using genomics to address questions in weed science [5, 18–20]. Training workshops taught commonly used bioinformatic pipelines, and oral and poster sessions showcased current research activities in weed genomics. At the conclusion of this meeting, attendees participated in a forward-looking discussion about the future of genomics in weed science and how the IWGC can help facilitate its successful implementation. In this paper, we summarize the goals of the IWGC and how we plan to provide support around the resources being developed to ensure they are widely accessible and utilized by the research community. We go on to highlight areas of research where these tools can be applied with hopes of attracting researchers from other fields to integrate weed science with the many other research areas where genomic tools are being successfully utilized, enabling new research towards adaptation, evolution, herbicide resistance, and genome biology.

### Development of Weed Genomics Resources by the IWGC

#### Reference Genomes and Data Analysis Tools

The first objective of the IWGC is to provide high quality genomic resources for agriculturally important weeds. The IWGC therefore created two main resources for information about, access to, or analysis of weed genomic data (Figure 1). The IWGC website [21] communicates the status and results of genome sequencing projects, information on training and funding opportunities, upcoming events, and news in weed genomics. It also contains details of all sequenced species including genome size, ploidy, chromosome number, herbicide resistance status, and reference genome assembly statistics. The IWGC either compiles existing data on genome size, ploidy, and chromosome number, or obtains the data using flow cytometry and cytogenetics (Figure 1; Additional File 1). Through this website, users can create an account to access our second main resource, an online genome database called WeedPedia. WeedPedia hosts IWGC-generated and other relevant publicly accessible genomic data as well as a suite of bioinformatic tools. Unlike what is available for other fields, weed science did not have a centralized hub for genomics information, data, and analysis prior to the IWGC. Our intention in creating WeedPedia is to encourage collaboration and equity of access to information across the research community. Importantly, all genome assemblies and annotations from the IWGC (Table 1), along with the raw data used to produce them, will be made available through NCBI GenBank. Upon completion of a one-year sponsoring member data confidentiality period for each species, scientific teams within the IWGC produce the first genome-wide investigation to submit for publication including whole genome level analyses on genes, gene families, and repetitive sequences, and comparative analysis with other species. Genome assemblies and data will be publicly available through NCBI as part of these initial publications for each species.

**Table 1.** Genome assemblies of 31 weed species completed or ongoing by the International Weed Genomics Consortium. All completed genomes are platinum assembly quality, defined as having chromosome-length scaffolds (i.e., 1-3 scaffolds per chromosome) for the assembly, unless indicated by *. Genome size estimated from flow cytometry or published references as indicated. + indicates that verification is currently in progress for cytogenetic information.

| Scientific name | Common name | Haplotypes in Assembly | Anticipated Availability Date | Ploidy | x | n | Genome Size Estimate (Gbp) |
| --- | --- | --- | --- | --- | --- | --- | --- |
| *Amaranthus hybridus* | smooth pigweed | 1;<br>Previous version [158] | November 2023 | diploid | 16 | 16 | 0.509 [159] |
| *Amaranthus palmeri* | Palmer amaranth | Previous version [158] | June 2024 | diploid | 17 | 17 | 0.445 [160] |
| *Amaranthus retroflexus* | redroot pigweed |  | In progress | diploid | 16 | 16 | 0.592 [160] |
| *Amaranthus tuberculatus* | common waterhemp | 2;<br>Previous version [158] | November 2023 | diploid | 16 | 16 | 0.694 [160] |
| *Ambrosia artemisiifolia* | common ragweed | Previous version [113] | In progress | diploid [161, 162] | 18 | 18 | 1.152 [163] |
| *Ambrosia trifida* | giant ragweed | 2 | December 2023 | diploid [161] | 12 | 12 | 1.872 [164] |
| *Apera spica-venti* | loose silkybent | 2 | November 2023 | diploid | 7 | 7 | 4.622 |
| *Avena fatua* | wild oat | 1 | November 2023 | hexaploid (Additional file 1) | 7 | 21 | 11.248 |
| *Chenopodium album* | common lambsquarters | 1 | November 2023 | hexaploid | 9 | 27 | 1.59 |
| *Cirsium arvense* | Canada thistle |  | In progress | diploid | 17 | 17 | 1.415 |
| *Convolvulus arvensis* | field bindweed |  | In progress | diploid<sup>+</sup> | 12<sup>+</sup> | 12<sup>+</sup> | 0.652 [163] |
| *Conyza bonariensis* (*Erigeron bonariensis*) | hairy fleabane |  | In progress | hexaploid [165] | 9 | 27 | 2.043 [166] |
| *Conyza sumatrensis* (*Erigeron sumatrensis*) | Sumatran fleabane | 1 | November 2023 | hexaploid | 9 | 27 | 1.874 |
| *Cyperus esculentus* | yellow nutsedge | 2 | December 2023 | diploid | 54 | 54 | 0.588 [167] |
| *Cyperus rotundus* | purple nutsedge | 2 | December 2023 | diploid | 54 | 54 | 0.49 [167] |
| *Digitaria insularis* | sourgrass | 1 | November 2023 | tetraploid | 9 | 18 | 1.529 |
| *Digitaria ischaemum* | hairy crabgrass |  | November 2024 | tetraploid | 9 | 18 | 0.625 |
| *Echinochloa colona* | junglerice (weedy genotype) | See crop genotype assembly by [89] | October 2024 | hexaploid | 9 | 27 | 1.372 [167] |
| *Euphorbia esula* | leafy spurge |  | In progress | hexaploid<sup>+</sup> [based on 168, 169] | 10<sup>+</sup> | 60<sup>+</sup> | 2.3 [170] |
| *Euphorbia heterophylla* | wild poinsettia |  | July 2024 | diploid [171] | 14 | 14 | Unknown, in progress |
| *Leptochloa chinensis* | Chinese sprangletop | 2;<br>See also [172] | November 2023 | tetraploid | 10 | 20 | 0.454 |
| *Lolium rigidum* | annual ryegrass | 2;<br>See also [173] | November 2023 | diploid (Additional file 1) | 7 | 7 | 2.41 |
| *Orobanche cernua* | nodding broomrape |  | In progress | diploid | 19 | 19 | 1.421 [174] |
| *Orobanche crenata* | crenate broomrape |  | In progress | diploid | 19 | 19 | 2.787 [174] |
| *Orobanche minor* | small broomrape |  | In progress | diploid | 19 | 19 | 1.792 [174] |
| *Parthenium hysterophorus* | ragweed parthenium |  | In progress | diploid [175] | 17 | 17 | Unknown, in progress |
| *Phalaris minor* | little seed canary grass | 1 | November 2023 | tetraploid (Additional file 1) | 7 | 14 | 5.851 |
| *\*Raphanus raphanistrum* | wild radish | Previous versions [176, 177] | In progress | diploid | 9 | 9 | 0.515 [176] |
| *Salsola tragus* | Russian thistle | 2 | November 2023 | tetraploid (Additional file 1) | 9 | 18 | 1.319 |
| *\*Sorghum halepense* | johnsongrass | 2 | January 2024 | tetraploid | 10 | 20 | 1.752 |
| *Verbascum blattaria* | moth mullein | 1 | December 2023 | diploid | 15 | 15 | 0.344 [178] |

WeedPedia is a cloud-based omics database management platform built from the software ‘CropPedia’, and licensed from KeyGene (Wageningen, The Netherlands). The interface allows users to access, visualize, and download genome assemblies along with structural and functional annotation. The platform includes a genome browser, comparative map viewer, pangenome tools, RNA-sequencing data visualization tools, genetic mapping and marker analysis tools, and alignment capabilities that allow searches by keyword or sequence. Additionally, genes encoding known target sites of herbicides have been specially annotated, allowing users to quickly identify and compare these genes of interest. The platform is flexible, making it compatible with future integration of other data types such as epigenetic or proteomic information. As an online platform with a graphical user interface, WeedPedia provides user-friendly, intuitive tools that encourage users to integrate genomics into their research. We aspire for WeedPedia to mimic the success of other public genomic databases such as NCBI, CoGe, Phytozome, InsectBase, and Mycocosm to name a few. WeedPedia currently hosts reference genomes for 40 species with additional genomes in the pipeline to reach a currently planned total of reference genomes for 53 species (Table 1). These genomes include both *de novo* reference genomes generated or in progress by the IWGC (31 species; Table 1), and publicly available genome assemblies of 22 weedy or related species (Table 2). As of October 2023, WeedPedia has over 300 registered users representing 27 countries spread across 6 continents.

**Table 2.**
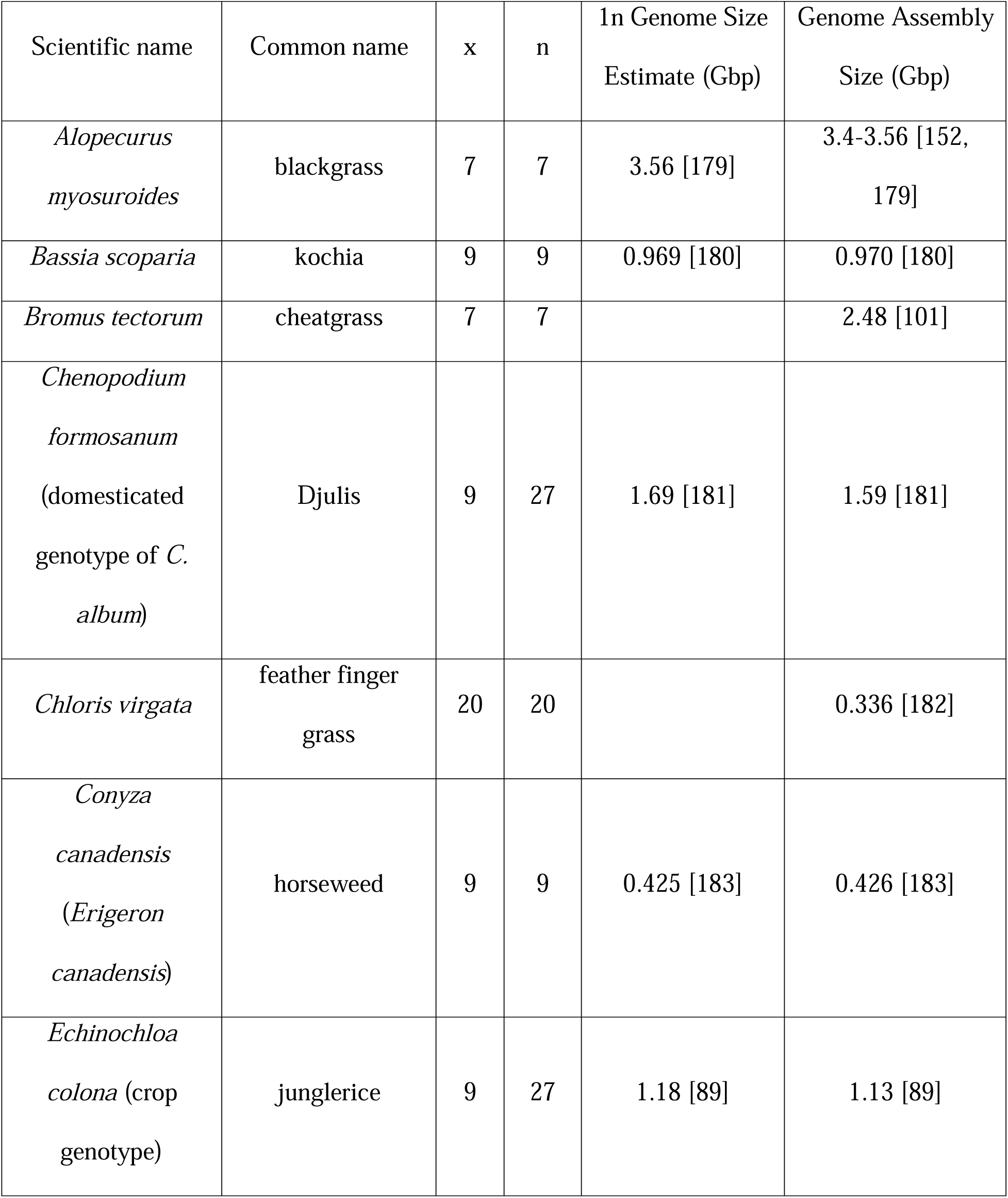
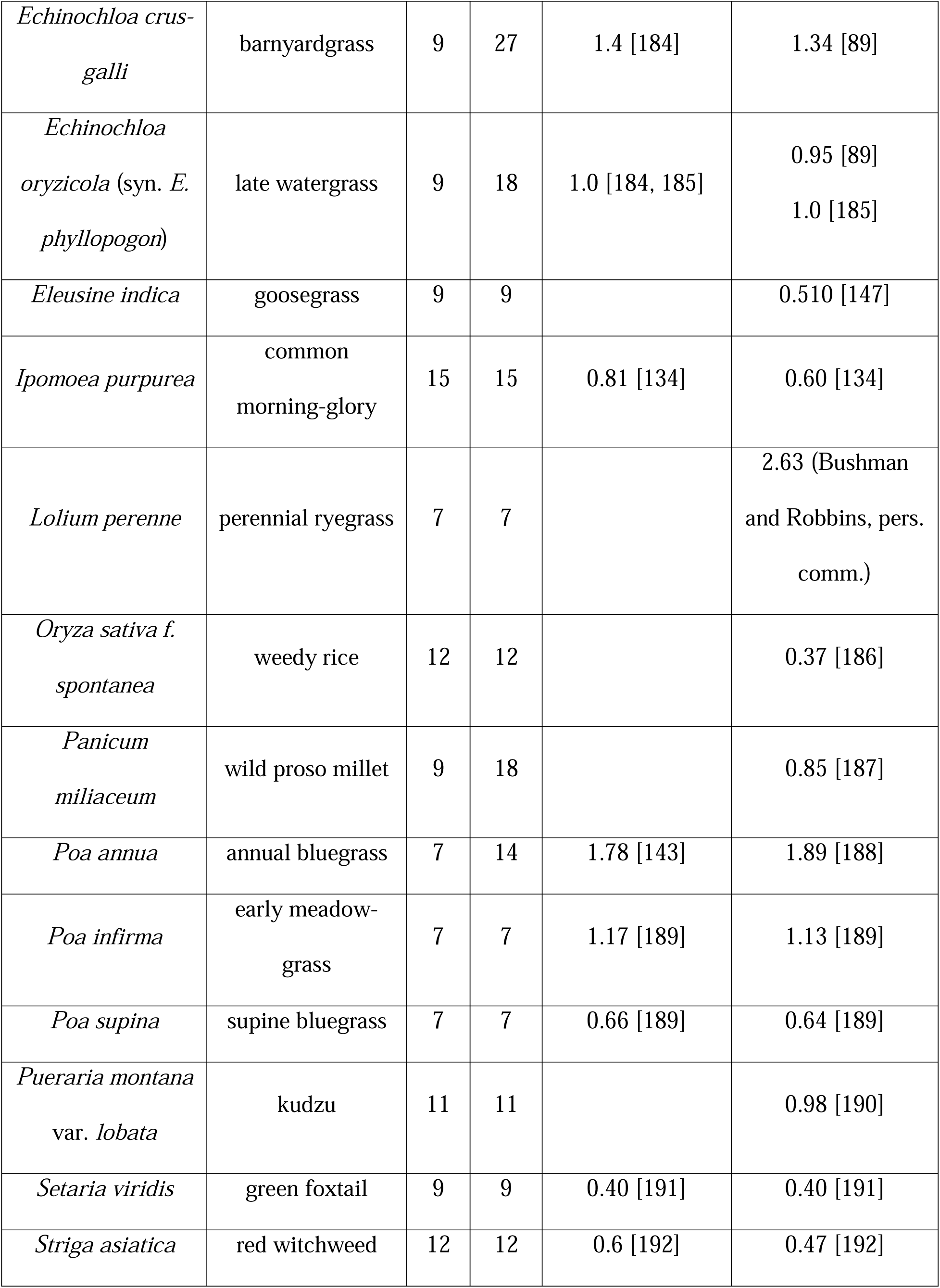
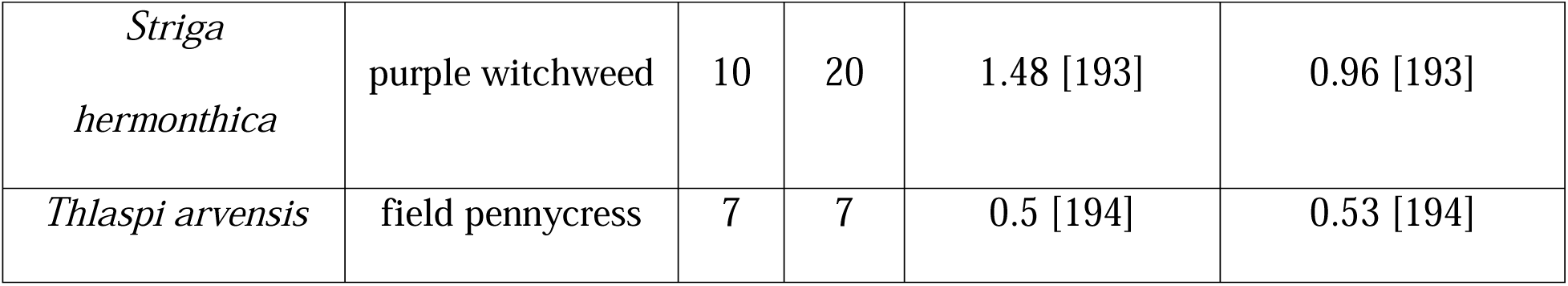
Genome assemblies and genomic information for 22 weed species produced by other groups independently of the International Weed Genomics Consortium. Haploid (1n) genome size estimations are either calculated through flow cytometry or k-mer estimation.

The IWGC reference genomes are generated in partnership with the Corteva Agriscience Genome Center of Excellence (Johnston, Iowa) using a combination of single molecule long read sequencing, optical genome maps, and chromosome conformation mapping. This strategy has yielded highly contiguous, phased, chromosome-level assemblies for 20 weed species, with additional assemblies currently in progress (Table 1). The IWGC assemblies have been completed as single or haplotype-resolved double-haplotype pseudomolecules in inbreeding and outbreeding species, respectively, with multiple genomes being near gapless. For example, the *de novo* assemblies of the allohexaploids *Conyza sumatrensis* and *Chenopodium album* have all chromosomes captured in single scaffolds and most chromosomes being gapless from telomere to telomere. Complementary full-length isoform (IsoSeq) sequencing of RNA collected from diverse tissue types and developmental stages assists in the development of gene models during annotation. Finally, the use of PacBio Revio has enabled the re-sequencing of 80 relevant accessions, which is enabling initial pangenomic analysis for some of the IWGC-selected species.

As with accessibility of data, a core objective of the IWGC is to facilitate open access to sequenced germplasm for all featured species. Historically, the weed science community has rarely shared or adopted standard germplasm (e.g., specific weed accessions). The IWGC has selected a specific accession of each species for reference genome assembly (typically susceptible to herbicides). In collaboration with a parallel effort by the Herbicide Resistant Plants committee of the Weed Science Society of America, seeds of the sequenced weed accessions will be deposited in the United States Department of Agriculture Germplasm Resources Information Network [22] for broad access by the scientific community. The IWGC ensures that sequenced accessions are collected and documented to comply with the Nagoya Protocol on access to genetic resources and the fair and equitable sharing of benefits arising from their utilization under the Convention on Biological Diversity and related Access and Benefit Sharing Legislation [23]. As additional accessions of weed species are sequenced (e.g., pangenomes are obtained) the IWGC will facilitate germplasm sharing protocols to support collaboration. Further, to simplify the investigation of herbicide resistance, the IWGC will link WeedPedia with the International Herbicide-Resistant Weed Database [24], an already widely known and utilized database for weed scientists.

#### Training and Collaboration in Weed Genomics

Beyond producing genomic tools and resources, a priority of the IWGC is to enable the utilization of these resources across a wide range of stakeholders. A holistic approach to training is required for weed science generally [25], and we would argue even more so for weed genomics. To accomplish our training goals, the IWGC is developing and delivering programs aimed at the full range of IWGC stakeholders and covering a breadth of relevant topics. We have taken care to ensure our approaches are diverse as to provide training to researchers with all levels of existing experience and differing reasons for engaging with these tools. Throughout, the focus is on ensuring that our training and outreach result in impacts that benefit a wide range of stakeholders.

Although recently developed tools are incredibly enabling and have great potential to replace antiquated methodology [26] and to solve pressing weed science problems [14], specialized computational skills are required to fully explore and unlock meaning from these highly complex datasets. Collaboration with, or training of, computational biologists equipped with these skills and resources developed by the IWGC will enable weed scientists to expand research programs and better understand the genetic underpinnings of weed evolution and herbicide resistance. To fill existing skill gaps, the IWGC is developing summer bootcamps and online modules directed specifically at weed scientists that will provide training on computational skills (Figure 1). Because successful utilization of the IWGC resources requires more than general computational skills, we have also created three additional targeted workshops that teach practical skills related to genomics databases, molecular biology, and population genomics (available at [27]).

Engagement opportunities during undergraduate degrees has been shown to improve academic outcomes [28, 29]. Therefore, the IWGC sponsors opportunities for undergraduates to undertake 10-week Research Experiences for Undergraduates (REU). These REU include an introduction to bioinformatics, a plant genomics research project that results in a presentation, and access to career building opportunities in diverse workplace environments. To increase equitable access to conferences and professional communities, we supported early career researchers to attend the first two IWGC conferences in the USA as well as workshops and bootcamps in Europe and South America. These hybrid or in-person travel grants are intentionally designed to remove barriers and increase participation of individuals from backgrounds and experiences currently underrepresented within weed/plant science or genomics [30]. Recipients of these travel awards gave presentations and gained the measurable benefits that come from either virtual or in-person participation in conferences [31]. Moving forward, weed scientists must amass skills associated with genomic analyses and collaborate with other area experts to fully leverage resources developed by the IWGC.

### Evolution of Weediness: Potential Research Utilizing New Weed Genomics Tools

Weeds can evolve from non-weed progenitors through wild colonization, crop de-domestication, or crop-wild hybridization [32]. Because the time span in which weeds have evolved is necessarily limited by the origins of agriculture, these non-weed relatives often still exist and can be leveraged through population genomic and comparative genomic approaches to identify the adaptive changes that have driven the evolution of weediness. The ability to rapidly adapt, persist, and spread in agroecosystems are defining features of weedy plants, leading many to advocate agricultural weeds as ideal candidates for studying rapid plant adaptation [33–36]. The insights gained from applying plant ecological approaches to the study of rapid weed adaptation will move us towards the ultimate goals of mitigating such adaptation and increasing the efficacy of crop breeding and biotechnology [14].

#### Biology and Ecological Genomics of Weeds

The impressive community effort to create and maintain resources for *Arabidopsis thaliana* ecological genomics provides a motivating example for the emerging study of weed genomics [37–40]. *Arabidopsis thaliana* was the first flowering plant species to have its genome fully sequenced [41] and rapidly became a model organism for plant molecular biology. As weedy genomes become available, collection, maintenance, and resequencing of globally distributed accessions of these species will help to replicate the success found in ecological studies of *A. thaliana* [42–48]. Evaluation of these accessions for traits of interest to produce large phenomics data sets (as in [49–53]) enables genome-wide association studies and population genomics analyses aimed at dissecting the genetic basis of variation in such traits [54]. Increasingly, these resources (e.g the 1001 genomes project [42]) have enabled *A. thaliana* to be utilized as a model species to explore the eco-evolutionary basis of plant adaptation in a more realistic ecological context. Weedy species should supplement lessons in eco-evolutionary genomics learned from these experiments in *A. thaliana*.

Untargeted genomic approaches for understanding the evolutionary trajectories of populations and the genetic basis of traits as described above rely on the collection of genotypic information from across the genome of many individuals. While whole-genome resequencing accomplishes this requirement and requires no custom methodology, this approach provides more information than is necessary and is prohibitively expensive in species with large genomes. Development and optimization of genotype-by-sequencing methods for capturing reduced representations of newly sequence genomes like those described by [55–57] will reduce the cost and computational requirements of genetic mapping and population genetic experiments. Additionally, the species sequenced by the IWGC do not currently have protocols for stable transformation, a key development in the popularity of *A. thaliana* as a model organism and a requirement for many functional genomic approaches. Functional validation of genes/variants believed to be responsible for traits of interest in weeds has thus far relied on transiently manipulating endogenous gene expression [58, 59] or ectopic expression of a transgene in a model system [60–62]. While these methods have been successful, few weed species have well-studied viral vectors to adapt for use in virus induced gene silencing and spray induced gene silencing is relatively ineffective without the use of nanocarriers [63], which require specialized equipment and expertise. Furthermore, traits with complex genetic architecture divergent between the researched and model species may not be amenable to functional genomic approaches using transgenesis techniques in model systems. Developing protocols for reduced representation sequencing, stable transformation, and gene editing/silencing in weeds will allow for more thorough characterization of candidate genetic variants underlying traits of interest.

Beyond rapid adaptation, some weedy species offer an opportunity to better understand co-evolution, like that between plants and pollinators and how their interaction leads to the spread of weedy alleles (Additional File 2). A suite of plant-insect traits has co-evolved to maximize the attraction of the insect pollinator community and the efficiency of pollen deposition between flowers ensuring fruit and seed production in many weeds [64, 65]. Genetic mapping experiments have identified genes and genetic variants responsible for many floral traits affecting pollinator interaction including petal color [66–69], flower symmetry and size [70–72], and production of volatile organic compounds [73–75] and nectar [76–78]. While these studies reveal candidate genes for selection under co-evolution, herbicide resistance alleles may also have pleiotropic effects on the ecology of weeds [79], altering plant-pollinator interactions [80]. Discovery of genes and genetic variants involved in weed-pollinator interaction and their molecular and environmental control may create opportunities for better management of weeds with insect-mediated pollination. For example, if management can disrupt pollinator attraction/interaction with these weeds, the efficiency of reproduction may be reduced.

A more complete understanding of weed ecological genomics will undoubtedly elucidate many unresolved questions regarding the genetic basis of various aspects of weediness. For instance, when comparing populations of a species from agricultural and non-agricultural environments, is there evidence for contemporary evolution of weedy traits selected by agricultural management or were ‘natural’ populations pre-adapted to agroecosystems? Where there is differentiation between weedy and natural populations, which traits are under selection and what is the genetic basis of variation in those traits? When comparing between weedy populations, is there evidence for parallel versus non-parallel evolution of weediness at the phenotypic and genotypic levels? Such studies may uncover fundamental truths about weediness. For example, is there a common phenotypic and/or genotypic basis for aspects of weediness amongst diverse weed species? As genomic tools developed by the IWGC enable researchers to address these questions, knowledge gained will help predict the potential development of newly important weed species in new environments and cropping systems.

#### Population Genomics

Weed species are certainly fierce competitors, able to outcompete crops and endemic species in their native environment, but they are also remarkable colonizers of perturbed habitats. Weeds achieve this through high fecundity, often producing tens of thousands of seeds per individual plant [81–83]. These large numbers in terms of demographic population size often combine with outcrossing reproduction to generate high levels of diversity with local effective population sizes in the hundreds of thousands [84, 85]. This has two important consequences: weed populations retain standing genetic variation and generate many new mutations, supporting weed success in the face of harsh control. The generation of genomic tools to monitor weed populations at the molecular level is a game-changer to understanding weed dynamics and precisely testing the effect of management on the genetic make-up of populations. We are now able to assess the effect of control on genetic diversity and monitor in real time whether these measures are effective at eroding weed populations.

Population genomic data, without any environmental or phenotypic information, can be used to scan the genomes of weed and non-weed relatives to identify selective sweeps, pointing at loci supporting weed adaptation on micro-or macro-evolutionary scales. Two recent within-species examples include weedy rice, where population differentiation between weedy and domesticated populations was used to identify the genetic basis of weedy de-domestication [86], and common waterhemp, where consistent allelic differences among natural and agricultural collections resolved a complex set of agriculturally adaptive alleles [87, 88]. A recent comparative population genomic study of weedy barnyardgrass and crop millet species has demonstrated how inter-specific investigations can resolve the signatures of crop and weed evolution [89] (also see [90] for a non-weed climate adaptation example). Multiple sequence alignments across numerous species provide complementary insight into adaptive convergence over deeper timescales, even with just one genomic sample per species (e.g., [91, 92]). Thus, the new IWGC weed genomes combined with genomes available for closely related crops (outlined by [14, 93]) and an effort to identify other non-weed wild relatives will be invaluable in characterizing the genetic architecture of weed adaptation and evolution across diverse species.

Weeds experience high levels of genetic selection, both artificial in response to agricultural practices and particularly herbicides, and natural in response to the environmental conditions they encounter [94, 95]. Using genomic analysis to identify loci that are the targets of selection and support adaptation would point at vulnerabilities that could be leveraged against weeds to develop new and more sustainable management strategies [96]. This is a key motivation to develop genotype-by-environment association (GEA) and selective sweep scan approaches, which allow researchers to resolve the molecular basis of multi-dimensional adaptation [97, 98]. GEA approaches, in particular, have been widely used on landscape-wide resequencing collections to determine the genetic basis of climate adaptation (e.g., [40, 99, 100]), but have yet to be fully exploited to diagnose the genetic basis of the various aspects of weediness [101]. Armed with data on environmental dimensions of agricultural settings, such as focal crop, soil quality, herbicide use, and climate, GEA approaches can help disentangle how discrete farming practices have influenced the evolution of weediness and resolve broader patterns of local adaptation across a weed’s range. Although non-weedy relatives are not technically required for GEA analyses, inclusion of environmental and genomic data from weed progenitors can further distinguish genetic variants underpinning weed origins from those involved in local adaptation.

New weeds emerge frequently [102], either through hybridization between species as documented for sea beet (*Beta vulgaris* ssp. *maritima)* hybridizing with crop beet to produce progeny that are well adapted to agricultural conditions [103–105], or through the invasion of alien species that find a new range to colonize. Biosecurity measures are often in place to stop the introduction of new weeds; however, the vast scale of global agricultural commodity trade precludes the possibility of total control. Population genomic analysis is now able to measure gene flow between populations [87, 106–108] and identify populations of origin for invasive species including weeds [109–111]. In particular, the invasion route of the pest fruitfly *Drosophila suzukii* from Eastern Asia to North America and Europe through Hawaii was deciphered using Approximate Bayesian Computation on high-throughput sequencing data from a global sample of multiple populations [112]. Genomics can also be leveraged to predict invasion rather than explain it. The resequencing of a global sample of common ragweed (*Ambrosia artemisiifolia* L.) elucidated a complex invasion route whereby Europe was invaded by multiple introductions of American ragweed that hybridized in Europe prior to a subsequent introduction to Australia [113, 114]. In this context, the use of genomically-informed species distribution models helps assess the risk associated with different source populations, which in the case of common ragweed, suggests that a source population from Florida would allow ragweed to invade most of northern Australia [115]. The IWGC, by federating research efforts globally, provides a platform that can support the transformation of biosecurity from perspective analysis towards predictive risk assessment.

#### Genetic Basis of Herbicide Resistance

Herbicide resistance is among the numerous weedy traits that can evolve in plant populations exposed to agricultural selection pressures. Over-reliance on herbicides to control weeds, along with low diversity and lack of redundancy in weed management strategies, has resulted in globally widespread herbicide resistance [116]. To date, 268 herbicide-resistant weed species have been reported worldwide, and at least one resistance case exists for 21 of the 31 existing herbicide sites of action [24] – significantly limiting chemical weed control options available to agriculturalists. This limitation of control options is exacerbated by the recent lack of discovery of herbicides with new sites of action [117].

Herbicide resistance may result from several different physiological mechanisms. Such mechanisms have been classified into two main groups, target-site resistance (TSR) [4, 118] and non-target-site resistance (NTSR) [4, 119]. The first group encompasses changes that reduce binding affinity between a herbicide and its target [120]. These changes may provide resistance to multiple herbicides that have a common biochemical target [121] and can be effectively managed through mixture and/or rotation of herbicides targeting different sites of action [122]. The second group (NTSR), includes alterations in herbicide absorption, translocation, sequestration, and/or metabolism that may lead to unpredictable pleotropic cross-resistance profiles where structurally and functionally diverse herbicides are rendered ineffective by one or more genetic variant(s) [60]. This mechanism of resistance threatens not only the efficacy of existing herbicidal chemistries, but also ones yet to be discovered. While TSR is well understood because of the ease of identification and molecular characterization of target site variants, NTSR mechanisms are significantly more challenging to research because they are often polygenic, and the resistance causing element(s) are not well understood [123].

Improving the current understanding of metabolic NTSR mechanisms is not an easy task, since genes of diverse biochemical functions are involved, many of which exist as extensive gene families [121, 124]. Expression changes of NTSR genes have been implicated in several resistance cases where the protein products of the genes are functionally equivalent across sensitive and resistant plants, but their relative abundance leads to resistance. Thus, regulatory elements of NTSR genes have been scrutinized to understand their role in NTSR mechanisms [125]. Similarly, epigenetic modifications have been hypothesized to play a role in NTSR, with much remaining to be explored [126–128]. Untargeted approaches such as genome-wide association, selective sweep scans, linkage mapping, RNA-sequencing, and metabolomic profiling have proven helpful to complement more specific biochemical- and chemo-characterization studies towards the elucidation of NTSR mechanisms as well as their regulation and evolution [60, 129–136]. Due to their complexity and importance, the IWGC has begun addressing this subject by manually curating the annotation of NTSR genes and developing a standard nomenclature for the gene families often involved in NTSR. This standardization will allow researchers to quickly identify true orthologous genes between weedy species, which is a hurdle for current research of these complex and often vast gene families.

High-quality weed genome assemblies and gene model annotations have helped and will be crucial for investigating the landscape of NTSR genes in weeds. They can also be used to predict the protein structure for herbicide target site and metabolism genes to predict the efficacy and selectivity of new candidate herbicides *in silico* to increase herbicide discovery throughput. Knowledge of the genetic basis of NTSR will aid the rational design of herbicides by 1) screening new compounds in the presence of newly discovered NTSR proteins during early research phases; 2) identifying conserved chemical structures that interact with these proteins; and 3) optimizing herbicide molecular design to lower potential for resistance evolution and increase potency/spectrum of control.

Moving forward, genomic resources will be increasingly needed and used not only for the design of conventional small molecule herbicides, but also for next generation technologies for sustainable weed management. Proteolysis targeting chimeras (PROTACs) have the potential to bind desired targets with great selectivity and degrade proteins by utilizing natural protein ubiquitination and degradation pathways within plants [137]. The combination of nanoparticles with oligonucleotides has recently shown potential to be used in spray applications towards gene silencing in weeds, which paves the way for a new, innovative, and sustainable method for weed management [138, 139]. Additionally, success in the field of pharmaceutical drug discovery in the development of molecules modulating protein-protein interactions offers another potential avenue towards the development of herbicides with novel targets [140, 141]. High-quality genomic references allow for the design of new weed management technologies like the ones listed here that are specific to – and effective across – weed species but have a null effect on non-target organisms. The tools being developed by the IWGC will have a crucial role in enabling the development of next generation weed management strategies that will reduce our reliance on the few chemical control options currently available to agriculturalists.

#### Comparative Genomics and Genome Biology

The genomes of weed species are as diverse as weed species themselves. Many weed species belong to unique plant families with no phylogenetically close model or crop species relatives for comparison. On all measurable metrics, weed genomes run the gamut. Some have smaller genomes like *Cyperus* spp. (∼0.26 Gb) while others are larger, such as *Avena fatua* (∼11.1 Gb) (Table 1). Some have high heterozygosity in terms of single nucleotide polymorphisms, repetitive DNA, and structural variants, such as the *Amaranthus* spp., while others are primarily self-pollinated and quite homozygous, such as *Poa annua* [142, 143]. Some are diploid such as *Conyza canadensis* and *Echinochloa haploclada* while others are polyploid such as *C. sumetrensis*, *E. crus-galli,* and *E. colona* [89]. The availability of genomic resources in these diverse, unexplored branches of the tree of life allows us to identify consistencies and anomalies in the field of genome biology.

The weed genomes published so far have focused mainly on weeds of agronomic crops, and studies have revolved around their ability to resist key herbicides. For example, genomic resources were vital in the elucidation of herbicide resistance cases involving target site gene copy number variants (CNVs). Gene CNVs of 5-enolpyruvylshikimate-3-phosphate synthase (*EPSPS*) have been found to confer resistance to the herbicide glyphosate in diverse weed species. To date, nine species have independently evolved *EPSPS* CNVs, and species achieve increased *EPSPS* copy number via different mechanisms [144]. For instance, the *EPSPS* CNV in *Bassia scoparia* is caused by tandem duplication, which is accredited to transposable element insertions flanking *EPSPS* and subsequent unequal crossing over events [145, 146]. In *Eleusine indica*, a *EPSPS* CNV was caused by translocation of the *EPSPS* locus into the subtelomere followed by telomeric sequence exchange [147]. One of the most fascinating genome biology discoveries in weed science has been that of extra-chromosomal circular DNAs (eccDNAs) that harbor the *EPSPS* gene in the weed species *Amaranthus palmeri* [148, 149]. In this case, the eccDNAs autonomously replicate separately from the nuclear genome and do not reintegrate into chromosomes, which has implications for inheritance, fitness, and genome structure [150]. These discoveries would not have been possible without reference assemblies of weed genomes, next-generation sequencing, and collaboration with experts in plant genomics and bioinformatics.

Another question that is often explored with weedy genomes is the nature and composition of gene families that are associated with NTSR. Gene families under consideration often include cytochrome P450s (CYPs), glutathione-*S*-transferases (GSTs), ABC transporters, etc. Some questions commonly considered with new weed genomes include: how many genes are in each of these gene families, where are they located, and which weed accessions and species have an over-abundance of them that might explain their ability to evolve resistance so rapidly [19, 89, 151, 152]? Weed genome resources are necessary to answer questions about gene family expansion or contraction during the evolution of weediness, including the role of polyploidy in NTSR gene family expansion as explored by [153].

#### Translational Research and Communication with Weed Management Stakeholders

Whereas genomics of model plants is typically aimed at addressing fundamental questions in plant biology, and genomics of crop species has the obvious goal of crop improvement, goals of genomics of weedy plants also include the development of more effective and sustainable strategies for their management. Weed genomics assists with these objectives by providing novel molecular ecological and evolutionary insights from the context of intensive anthropogenic management (which is lacking in model plants), and offers knowledge and resources for trait discovery for crop improvement, especially given that many wild crop relatives are also important agronomic weeds (e.g. [154]). For instance, crop-wild relatives are valuable for improving crop breeding for marginal environments [155]. Thus, weed genomics presents unique opportunities and challenges relative to plant genomics more broadly. It should also be noted that although weed science at its core is a very applied discipline, it draws broadly from many scientific disciplines such as, plant physiology, chemistry, ecology, and evolutionary biology, to name a few. The successful integration of weed-management strategies, therefore, requires extensive collaboration among individuals collectively possessing the necessary expertise [156]. Consequently, a major objective of the IWGC is to ensure that basic findings arising from weed genomics are translated to advances in weed management and crop breeding by collaborating broadly with breeders, applied weed scientists, outreach specialists, and practitioners.

To accomplish this objective, the IWGC must facilitate communication of weed genomics findings to relevant stakeholders (Figure 1). With the growing complexity of herbicide resistance management, practitioners are beginning to recognize the importance of understanding resistance mechanisms to inform appropriate management tactics [14]. Although weed science practitioners do not need to understand the technical details of weed genomics, their appreciation of the power of weed genomics-together with their unique insights from field observations - will yield novel opportunities for applications of weed genomics to weed management. In particular, combining field management history with information on weed resistance mechanisms is expected to provide novel insights into evolutionary trajectories [e.g., 6, 157], which can be utilized for disrupting evolutionary adaptation. It can be difficult to obtain field history information from practitioners, but developing an understanding among them of the importance of such information can be invaluable. To address these aspects, the IWGC can provide funding, or at least coordinate teams, to build extension/education programs focused on weed genomics. Factsheets and easy-to-understand infographics can be developed and communicated to various stakeholders through traditional and electronic media.

## Conclusions

Weeds are unique and fascinating plants, having significant impacts on agriculture and ecosystems; and yet, aspects of their biology, ecology, and genetics remain poorly understood. Weeds represent a unique area within plant biology, given their repeated rapid adaptation to sudden and severe shifts in the selective landscape of anthropogenic management practices. The production of a public genomics database with reference genomes for over 50 weed species represents a substantial step forward towards research goals that improve our understanding of the biology and evolution of weeds. Future work is needed to improve annotations, particularly for complex gene families involved in herbicide detoxification, structural variants, and mobile genetic elements, given the evidence to date of the generation of adaptive genetic variation in weeds through structural variation. As reference genome assemblies become available; standard, affordable methods for gathering genotype information will allow for the identification of genetic variants underlying traits of interest. Further, development of methods for functional gene validation and hypothesis testing is needed in weeds to validate the effect of genetic variants detected through such experiments, including systems for transformation, gene editing, and transient gene silencing and expression. Future research should focus on utilizing weed genomes to investigate questions about the evolutionary biology, ecology, and genetics of weedy traits and weed population dynamics. The IWGC plans to continue the public-private partnership model to continue to host the WeedPedia database, integrate new datasets such as genome resequencing and transcriptomes, conduct trainings, and serve as a research coordination network to ensure that advances in weed science from around the world are shared across the research community (Figure 1). Bridging basic plant genomics with translational applications in weeds is needed to deliver on the potential of weed genomics to improve weed management and crop breeding.

## Availability of data and materials

The datasets supporting the conclusions of this article is included within the article and its additional files. All genome assemblies and related sequencing data produced by the IWGC will be available through NCBI as part of publications reporting the first genome-wide analysis for each species.

## Acknowledgements

The International Weed Genomics Consortium is supported by BASF SE, Bayer AG, Syngenta Ltd, Corteva Agriscience, CropLife International, the Foundation for Food and Agriculture Research (Award DSnew-024), and a conference grant from USDA-NIFA (Award number 2021-67013-33570).

## Additional Files

Additional File 1 (.docx). Methods and results for visualizing and counting the metaphase chromosomes of (1A): diploid *Lolium rigidum*; (1B): hexaploid *Avena fatua*; (1C): diploid *Phalaris minor*; and (1D): tetraploid *Salsola tragus*.

Additional File 2 (.docx). List of completed and in-progress genome assemblies of weed species pollinated by insects.

