## Additional File 1 for "The International Weed Genomics Consortium: Community Resources for Weed Genomics Research"

Additional File 1. Methods and results for visualizing and counting the metaphase chromosomes of: diploid *Lolium rigidum*; hexaploid *Avena fatua*; diploid *Phalaris minor*; and tetraploid *Salsola tragus*.

**Preparation of metaphase chromosomes:**

Young actively growing root tips were collected early in the morning (before 8 am) and treated with $N_2O$ gas at 10 bar for 3 h, followed by a pre-treatment with α-bromonapthalene for 8 h at 4 °C, and finally, overnight fixation in 3:1 (ethanol: acetic acid) solution at room temperature. The root tips were incubated in a humid chamber with an enzyme mixture containing 0.3% pectolyase, 0.2% cellulase, and 0.2% cytohelicase for 45 min at 37 °C. To acquire well-spread metaphase chromosomes, the macerated root tips were split and squashed with 65% acetic acid (20 µl/slide). The slides were subsequently stored at -80 °C until further use. Vectashield solution diluted with 2% 4, 6-diamidino-2-phenylindole (DAPI) was used to observe metaphase chromosomes under the microscope. All the images were captured at 60× emulsion oil magnification using an Olympus BX61 fluorescent microscope equipped with a Hamamatsu C10600 camera and further processed with the MetaMorph software.

### Annual ryegrass *(Lolium rigidum)*

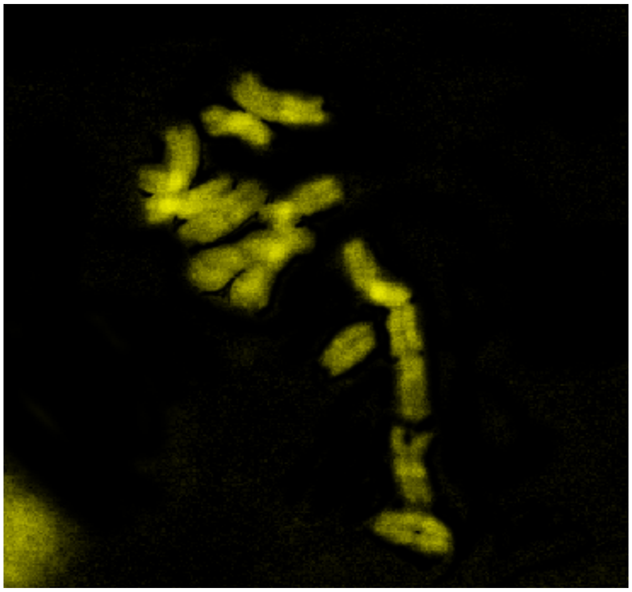

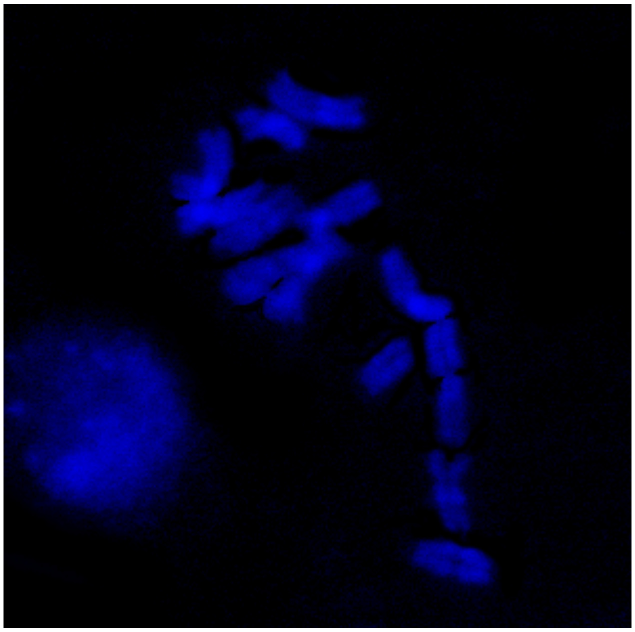

$2n = 2x = 14$

### Wild Oat (*Avena fatua*)

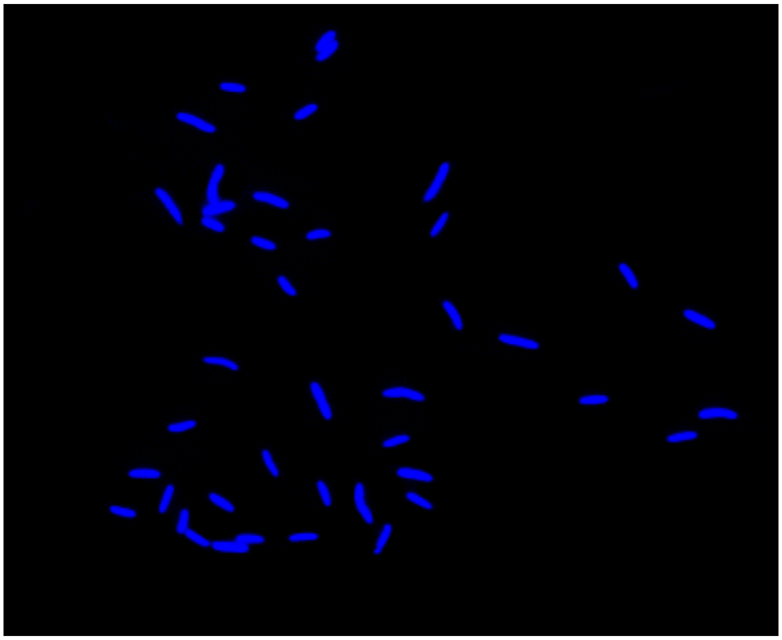

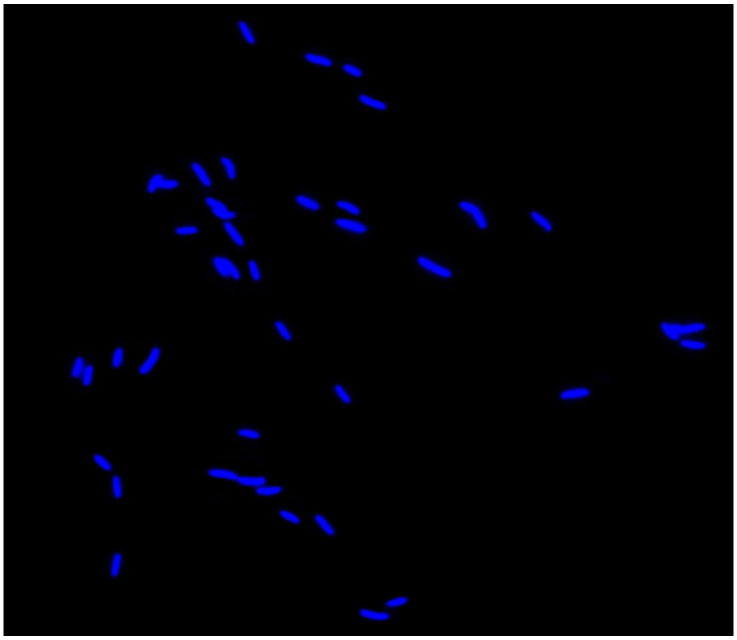

$2n = 6x = 42$

**Little seed canary grass**
***(Phalaris minor)***

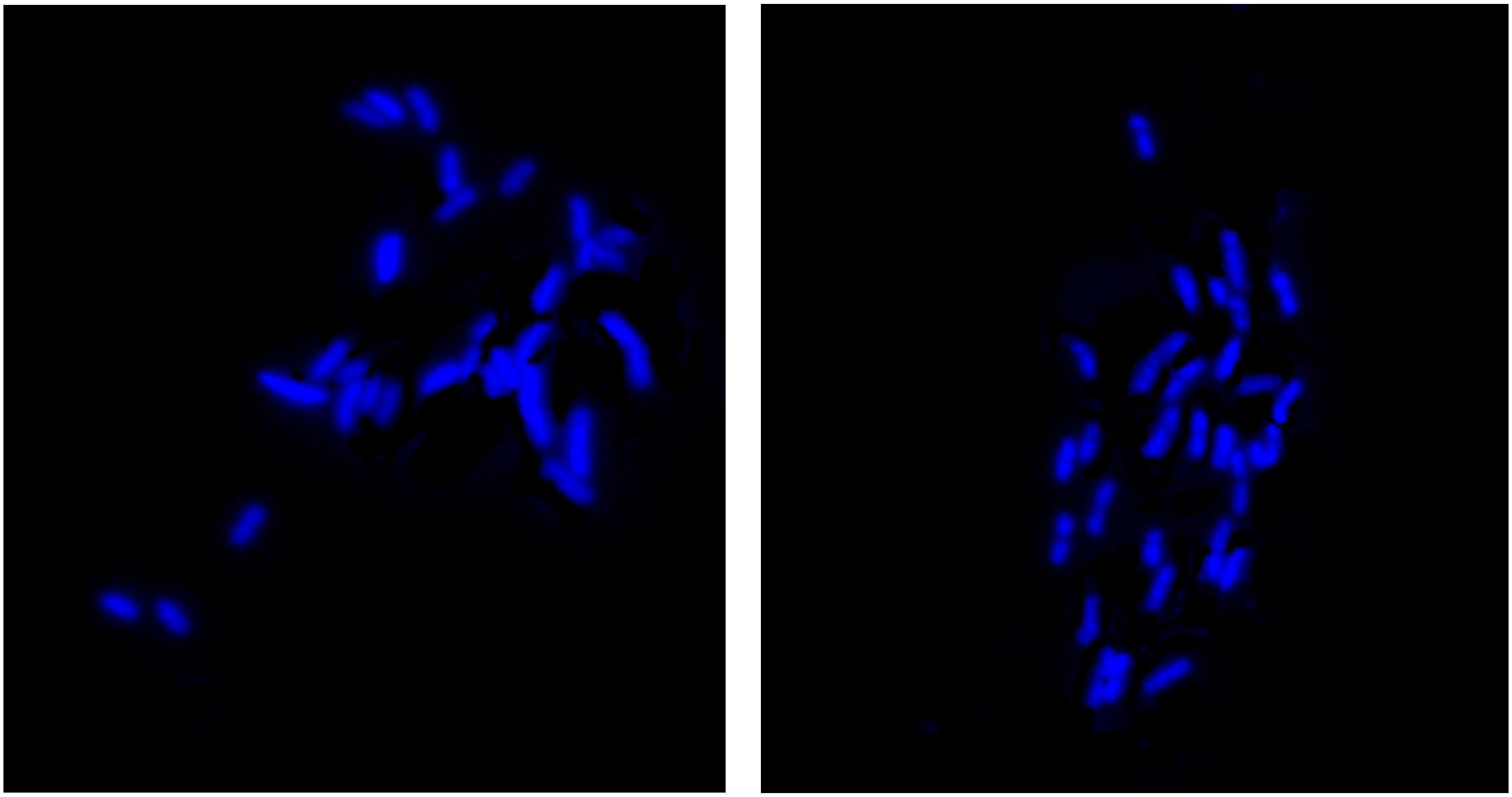

2n =2x = 28

### Russian thistle (*Salsola tragus*)

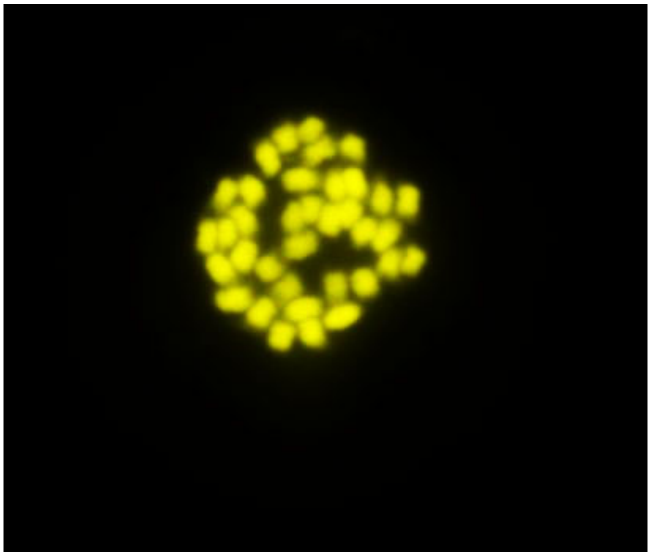

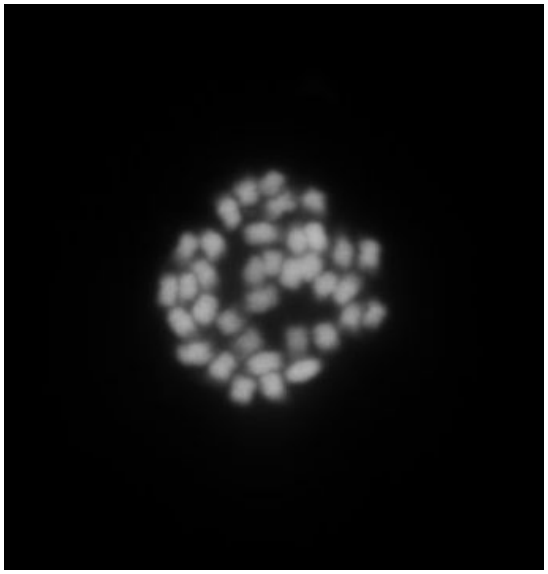

$2n = 4x = 36$
