## Additional File 2 for "The International Weed Genomics Consortium: Community Resources for Weed Genomics Research"

Additional File 2: Completed and in-progress genome assemblies of weed species pollinated by insects.

| **Weed species** | **Family** | **Entomophilous pollination** | **Reference** |
|---|---|---|---|
| *Cirsium arvense* | Asteraceae | Yes | [1] |
| *Conyza canadensis* | Asteraceae | Primarily self-pollinating, although insects were observed visiting flowers | [2] |
| *Erigeron sumatrensis* | Asteraceae | Yes | [3] |
| *Parthenium hysterophorus* | Asteraceae | Yes | [4, 5] |
| *Raphanus raphanistrum* | Brassicaceae | Yes | [6, 7] |
| *Salsola tragus* | Chenopodiaceae | Mainly anemophilous but also visited by insects | [8] |
| *Convolvulus arvensis* | Convolvulaceae | Yes | [9] |
| *Ipomoea purpurea* | Convolvulaceae | Yes | [10] |
| *Euphorbia esula* | Euphorbiaceae | Yes | [11] |
| *Euphorbia heterophylla* | Euphorbiaceae | Yes | [12] |
| *Verbascum blattaria* | Scrophulariaceae | Yes | [13] |
